# Susceptibility status of larval *Aedes aegypti* mosquitoes in Saudi Arabia

**DOI:** 10.1101/2020.12.14.422731

**Authors:** Ashwaq M Al Nazawi, Simon Ashall, David Weetman

**Affiliations:** Preventive Medicine Department, Public Health Directorate, Ministry of Health, Jeddah 22246, Saudi Arabia; School of Life Sciences, Keele University, Staffordshire, ST5 5BG, UK; Department of Vector Biology, Liverpool School of Tropical Medicine, Pembroke Place, Liverpool L3 5QA, UK

**Keywords:** Mosquito larvae, Larval bioassay, *Bti*, temephos

## Abstract

Vector control programs worldwide are facing the challenge of mosquitoes becoming resistant to available insecticides. Larviciding is a crucial preventative measure for dengue control but data on insecticide resistance of larval *Ae. aegypti* in the Middle Eastern Region are limited. This study assesses the susceptibility status of *Ae. aegypti* collected from the two most important dengue foci in Saudi Arabia, Jeddah and Makkah, to important chemical and biological larvicides; the organophosphate temephos and *Bacillus thuringiensis israelensis*, *Bti*). Whilst worldwide, and particularly in Latin America, high-level resistance to temephos is common, Jeddah and Makkah populations exhibited full susceptibility to both temephos and *Bti.* These data suggest each can be considered by vector control programs for preventative dengue control in the region, as part of temporal rotations or spatial mosaics to manage insecticide resistance.

## 1. Introduction

In Saudi Arabia, insecticides are extensively used to combat mosquito-borne diseases and other household pests, as well as in agriculture (1). *Aedes aegypti* is primarily controlled by larvicides such as Spinosad (Natular®), *Bacillus thuringiensis israelensis* (*Bti*) toxin (VectoBac**®**), pyriproxyfen and diflubenzuron. Adulticides such as deltamethrin, permethrin, cyfluthrin and fenitrothion are also used for fogging and indoor residual spraying to reduce the density of adult mosquitoes during outbreak situations (2). Temephos, *Bti*, Spinosad and insect growth regulatory hormones such as pyriproxyfen are used as larvicides in breeding sites, but *Bti* and Spinosad are more common in Jeddah and Makkah (1, 3, 4). However, the extensive use of chemical insecticides has led to the development of insecticide resistance in *Ae. aegypti* worldwide including Saudi Arabia (4–11). In 2011*, Ae. aegypti* strains from Makkah were found to be resistant to lambda-cyhalothrin, deltamethrin, permethrin, bendiocarb and cyfluthrin (10, 11) but still susceptible to pirimiphos-methyl (actellic) and *Bacillus thuringiensis israelensis Bti* (Bacilod) (11). In addition, Jeddah strains showed high prevalence of resistance to the pyrethroid deltamethrin and permethrin and the carbamate bendiocarb (10) but no studies to date have been considered larvicides. In Jazan, the population was resistant to lambda-cyhalothrin, DDT, bendiocarb and showed moderate resistance to permethrin, deltamethrin and fenitrothion (yet remained susceptible to cyfluthrin) (4). The larvae were reported as highly resistant to temephos, but the documented LC_50_ of 61.8 mg/L, appears unfeasibly high being far beyond the LC_50_ reported for other temephos resistant populations in the world (12, 13) suggesting further investigation is essential.

A major limitation of the control program in the region is the limited surveillance to monitor the effectiveness of control intervention, or changes in the resistance of populations that may undermine the control efforts. We therefore assess the susceptibility status of the sole local dengue vector *Ae. aegypti* collected from Makkah of Saudi Arabia to larvicides (temephos and *Bti*). The outcome of this study will provide reliable, updated data on the resistance profile of larval *Ae. aegypti* populations from Saudi Arabia and may provide indication of which insecticides may be more effective.

## 2. Materials and Methods

### 2.1 Mosquito strain

*Aedes aegypti* larvae were collected from multiple breeding sites in two dengue endemic areas in Makkah (Lab= 21°45’2.13 N, 39°92’1.96 E; field=21°40’7.70 N, 39°86’3.19 E) and Jeddah (Lab=21°35’2.13 N, 39°13’9.42 E; field=21°60’3.97 N, 39°27’2.49 E). The lab strains were fifth generation from the original field which was collected in 03-04/2016. The field strain was collected in 01-02/2018. The larvae were reared as described by (10). Two reference strains, Cayman, a multiply resistant lab strain, though reported as lacking temephos resistance (14), and the standard (ubiquitously-susceptible) strain New Orleans were used. All strains were raised under the same standard insectary conditions at the Liverpool School of Tropical Medicine (10).

### 2.2 Larval Bioassays

Larval bioassays were carried out on *Aedes* strains shown in **Table 1** according to the WHO protocol (15) to determine the lethal concentrations (LC_50_) and the resistance ratio relative to New Orleans (RR_50_).

**Table 1.** Number of *Ae. aegypti* larvae from Jeddah and Makkah used in each bioassay at different times.

| Temephos assay |  | *Bacillus thuringiensis israelensis* assay |  |
| --- | --- | --- | --- |
| Strain | Sample size | Strain | Sample size |
| New Orleans | 900 | New Orleans | 650 |
| Makkah lab strain | 900 | Makkah lab strain | 590 |
| Jeddah lab strain | 900 | Jeddah lab strain | 650 |
|  |  | Makkah field strain | 546 |
|  |  | Jeddah field strain | 567 |
|  |  | Cayman | 605 |

Bioassays were performed using temephos (Sigma-Aldrich, Dorset, UK), or *Bti* (Vectobac®12AS 1.2%,1200 ITU/mg. A total of eight different concentrations of *Bti* and nine of temephos were used for each strain **(Table 2&3)**. The concentrations were selected as they have been reported to result in larval mortality between 10% and 95% (15). The data was used to calculate the lethal dose that kills 50% (LC_50_) in each population. Dilutions of temephos (stock dissolved in absolute ethanol) with distilled water up to a total volume of 200mL are detailed in **Table 2.** For each concentration of each insecticide, three or four replicates of a pool of approximately 25 late third or early fourth instar larvae were tested along with a negative control pool;1mL absolute ethanol mixed into 199 mL of distilled water for temephos or into 100ml of distilled water for *Bti* assays.

**Table 2.** Temephos stock dilution with distilled water up to 200 ml to obtain the appropriate final concentrations.

| Final concentration (mg/L) | Volume of stock solution (mL) | Volume of distilled water (mL) |
| --- | --- | --- |
| 0.08 | 1 | 199 |
| 0.07 | 0.875 | 199.125 |
| 0.06 | 0.75 | 199.25 |
| 0.04 | 0.5 | 199.5 |
| 0.03 | 0.375 | 199.625 |
| 0.02 | 0.25 | 199.75 |
| 0.01 | 0.125 | 199.875 |
| 0.005 | 0.0625 | 199.9375 |
| 0.0025 | 0.03125 | 199.96875 |

**Table 3.** *Bacillus thuringiensis israelensis (Bti)* stock dilution with distilled water from the stock solution at 1.2%.

| C1 (stock) ppm | Volume of stock solution (mL) | Desired concentration (100mL)-ppm | Volume 2 (mL) |
| --- | --- | --- | --- |
| 120 | 0.005 | 0.0059997 | 100.005 |
| 120 | 0.003 | 0.003599892 | 100.003 |
| 120 | 0.002 | 0.002399952 | 100.002 |
| 120 | 0.001 | 0.001199988 | 100.001 |
| 120 | 0.00075 | 0.000899993 | 100.00075 |
| 120 | 0.0005 | 0.000599997 | 100.0005 |
| 120 | 0.0002 | 0.00024 | 100.0002 |
| 120 | 0.0001 | 0.00012 | 100.0001 |

Vectobac stock (1.2%) was diluted by adding 1ml of the stock (1.2%) to 99 ml distilled water to obtain 0.012% (120ppm) which was used in the experiment.

All larval bioassays were performed in 6 cm in diameter plastic bowls; Mortality was recorded after 24h of exposure. Any larvae failing or unable to swim up to the surface independently were counted as dead. Any larvae that had pupated during exposure were omitted from the total count.

### 2.3 Statistical analysis

The mortality (%) was calculated for the number of mosquitoes or larvae that were dead after 24h exposure. The LC_50_ value for the larval bioassays was calculated using probit regression analysis (SPSS version 24). If chisq >0.05, confidence limits were adjusted accordingly (SPSS does this unless the fit is terrible, if it is it will not calculated Cis). The resistance ratio (RR) was calculated by comparison of the resistant Makkah and Jeddah strains against the susceptible New Orleans strain using the formula below to monitor the level of insecticide resistance in a field population.

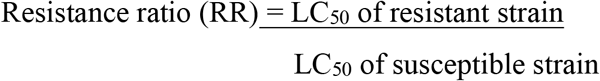

## 3. Results & Discussion

### 3.1 Larval bioassays

Mortality was not observed in any strain in the control assays. Based on the mortality rate across different concentrations of temephos and *Bti*, resistance to the larvicides was higher in field strains when compared to the New Orleans strain (**Table 4 and Table 5**).

**Table 4.** Average percentage mortality of *Ae. aegypti* larvae from Makkah and Jeddah and the susceptible strain, New Orleans exposed to nine concentrations of temephos.

| Strain | Temephos Concentration (mg/L) |  |  |  |  |  |  |  |  |
| --- | --- | --- | --- | --- | --- | --- | --- | --- | --- |
|  | 0.0025 (%) | 0.005 (%) | 0.01 (%) | 0.02 (%) | 0.03 (%) | 0.04 (%) | 0.06 (%) | 0.07 (%) | 0.08 (%) |
| Makkah | 0 | 0 | 9.18 | 70 | 89.6 | 100 | 99 | 99 | 100 |
| Jeddah | 1 | 0 | 1.02 | 32 | 79 | 73 | 94.8 | 98 | 100 |
| New Orleans | 5.2 | 9.9 | 52.6 | 81 | 96 | 99 | 100 | 100 | 100 |

**Table 5.**
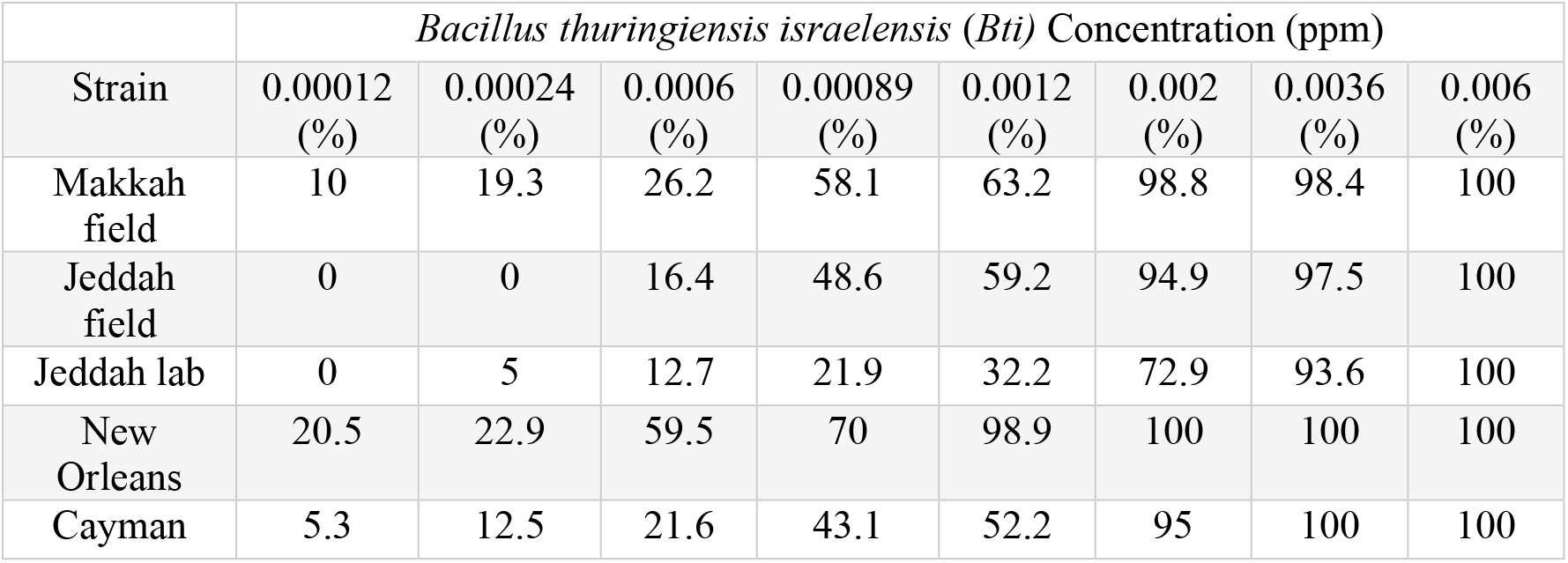
Average percentage mortality of *Ae. aegypti* larvae from Makkah, Jeddah (field and lab strains), New Orleans and Cayman strains exposed to eight concentrations of *Bti*.

| Strain | *Bacillus thuringiensis israelensis* (*Bti*) Concentration (ppm) |  |  |  |  |  |  |  |
| --- | --- | --- | --- | --- | --- | --- | --- | --- |
|  | 0.00012<br>(%) | 0.00024<br>(%) | 0.0006<br>(%) | 0.00089<br>(%) | 0.0012<br>(%) | 0.002<br>(%) | 0.0036<br>(%) | 0.006<br>(%) |
| Makkah field | 10 | 19.3 | 26.2 | 58.1 | 63.2 | 98.8 | 98.4 | 100 |
| Jeddah field | 0 | 0 | 16.4 | 48.6 | 59.2 | 94.9 | 97.5 | 100 |
| Jeddah lab | 0 | 5 | 12.7 | 21.9 | 32.2 | 72.9 | 93.6 | 100 |
| New Orleans | 20.5 | 22.9 | 59.5 | 70 | 98.9 | 100 | 100 | 100 |
| Cayman | 5.3 | 12.5 | 21.6 | 43.1 | 52.2 | 95 | 100 | 100 |

Indeed in both the temephos bioassays, the LC_50_ confidence intervals were not overlapping in comparisons of any strain, indicating a significant difference in mortality between the strains (**Table 6)**. However, whilst there is significant variation in susceptibility, current guidelines (16), suggest that a resistance ratio <5 indicates limited/no resistance; 5-10 moderate resistance, and >10 is substantial resistance. Therefore, based on this classification, no definitive resistance to temephos and *Bti* was identified in any of the strains tested.

**Table 6.** Lethal concentrations of temephos and *Bti* that kills 50% of *Ae. aegypti* strains.

| Strain | Temephos assay |  | *Bti* assay |  |
| --- | --- | --- | --- | --- |
|  | LC<sub>50</sub>, mg/L<br>(95% C.I.) | RR | LC<sub>50</sub>, ppm<br>(95% C.I.) | RR |
| New Orleans | 0.010<sup>a</sup><br>(0.009-0.011) | 1 | 0.00041<sup>a</sup><br>(0.000276-0.000537) | 1 |
| Makkah lab | 0.017<sup>b</sup><br>(0.014-0.019) | 1.7 | n/a | n/a |
| Jeddah lab | 0.029<sup>c</sup><br>(0.025-0.034) | 2.9 | 0.001483<sup>b</sup><br>(0.001341-0.001629) | 3.6 |
| Makkah field | n/a |  | 0.000834<sup>c</sup><br>(0.000688-0.000982) | 2.1 |
| Jeddah field | n/a |  | 0.00098<sup>b,c</sup><br>(0.000882-0.001076) | 2.4 |
| Cayman | n/a |  | 0.001018<sup>b,c</sup><br>(0.000882-0.001157) | 2.5 |

Shared letters within a column indicate no significant difference based on overlapping confidence limits. The Makkah lab *Bti* assay was not calculated (n/c) because a very poor fit of the probit model meant that LC<sub>50</sub> confidence intervals could not be reliably estimated even with a correction factor. Makkah and Jeddah field strains were not assessed (n/a) with temephos because the strains were not available at LSTM. The RR shown in the table indicates the resistance ratio.

The current study was conducted to assess the susceptibility of larval *Ae. aegypti* to commonly used insecticides in the cities of Jeddah and Makkah. Larval bioassays did not detect resistance in either Makkah or Jeddah to temephos or *Bti* (all resistance ratios <5 compared to a standard susceptible strain (17). In contrast, extreme temephos resistance in *Ae. aegypti* larvae from Jazan (LC_50_=61.8 mg/L) was reported in 2016 (4). When compared to the average LC_50_ of multiple separate studies of the susceptible reference strains Rockefeller, New Orleans and Bora Bora (18), this equates to a resistance ratio above 10,000, far exceeding the highest ratio of 224 previously recorded (in Brazil; (18)). This estimate from Jazan thus appears unlikely to be correct, and in the absence of additional data, a provisional assessment of temephos susceptibility in Saudi Arabia seems appropriate.

Temephos resistance in *Ae. aegypti* larvae has been recorded globally including British Virgin Islands (19), Thailand (20), Brazil (21), Cuba (22), Colombia (23), Martinique (24) and Santiago island (25). Whilst the current data suggest susceptibility, it is important to note that both Saudi Arabian strains showed significantly higher LC_50_ values than the susceptible New Orleans strain.

*Bacillus thuringiensis israelensis* (*Bti*) is a bacterial derived toxin that has been widely used for vector control. The populations from Jeddah and Makkah were susceptible to this compound in comparison (based on a resistance ratio <5) with the New Orleans strain. Almost all other studies have reported similar findings, including Martinique populations (highly resistant to most insecticides) that were susceptible to *Bti* compared to the Bora-Bora strain, Santiago island, Cameroon and Malaysia (18). Although *Bti* resistance is apparently absent in *Ae. aegypti* populations to date, resistance has detected in *Culex pipiens,* from Syracuse, New York which had a resistance ratio of 33-fold when compared to the S-Lab susceptible strain (26). Resistance to *Bti* has also been demonstrated in *Aedes rusticus Rossi* mosquitoes, selected for resistance through annual *Bti* treatment in larval sites in the Rhône-Alpes region. The mosquitoes collected in the treatment area had a moderate resistance ratio up to 7.9 fold compared to the untreated area (27). The attained resistance levels were still relatively low compared to when mosquitoes are selected for resistance to other insecticides (27). The multiple active toxins-Cry4A, Cry4B, Cry11A and Cyt1A-produced by *Bti* might act at different receptors, making evolution of resistance to *Bti* very difficult (28). Nevertheless, both Saudi populations showed a significantly higher LC_50_ value than the New Orleans strain, suggesting that, as with temephos, variation exists which might be selected to higher levels in future.

Therefore, although the *Ae. aegypti* population in Jeddah and Makkah are still susceptible to temephos, rotational application of *Bti* and temephos, or another larvicide to which there is full susceptibility, would be advisable to slow down evolution of resistance to either of them thus retaining their efficacy over extended periods of use in vector control. Our findings suggest the potential to develop resistance to both insecticides may exist and thus mixture or rotation is advisable, along with continued monitoring, and consideration of other options such as insecticide growth regulators.

## 4. Conclusion

*Aedes aegypti* from Makkah and Jeddah remain susceptible to the larvicides assessed in this study and thus larval source management and larviciding could remain an effective tool in control, but regular monitoring and consideration of additional alternatives is advised.

## Acknowledgment

The authors are indebted all the municipal staff (administrators and laboratory technicians in the vector control programme) of Makkah and Jeddah in Saudi Arabia. We are also thankful to Dr Craig Wilding in Liverpool John Moores University for his co-operation during this work.

## Notes

### Competing Interest Statement

The authors have declared no competing interest.

